# Cell fate specification and conversion generate foveolar cone subtype patterning in human retinal organoids

**DOI:** 10.1101/2023.01.28.526051

**Authors:** Katarzyna A. Hussey, Kiara Eldred, Brian Guy, Clayton Santiago, Ian Glass, Thomas A. Reh, Seth Blackshaw, Loyal A. Goff, Robert J. Johnston

## Abstract

The mechanisms that generate patterns of cell types unique to humans are poorly understood. In the central region of the human retina, the high-acuity foveola is notable, in part, for its dense packing of green (M) and red (L) cones and absence of blue (S) cones. To identify mechanisms that promote M/L and suppress S cone patterning in the foveola, we examined human fetal retinas and differentiated human retinal organoids. During development, sparse S-opsin-expressing cones are initially observed in the foveola. Later in fetal development, the foveola contains a mix of cones that either co-express S- and M/L-opsins or exclusively express M/L-opsin. In adults, only M/L cones are present. Two signaling pathway regulators are highly and continuously expressed in the central retina: Cytochrome P450 26 subfamily A member 1 (CYP26A1), which degrades retinoic acid (RA) and Deiodinase 2 (DIO2), which promotes thyroid hormone (TH) signaling. Both *CYP26A1* null mutant organoids and high RA conditions increased the number of S cones and reduced the number of M/L cones in human retinal organoids. In contrast, sustained TH signaling promoted the generation of M/L-opsin-expressing cones and induced M/L-opsin expression in S-opsin-expressing cones, showing that cone fate is plastic. Our data suggest that CYP26A1 degrades RA to specify M/L cones and limit S cones and that continuous DIO2 expression sustains high levels of TH to convert S cones into M/L cones, resulting in the foveola containing only M/L cones. Since the foveola is highly susceptible to impairment in diseases such as macular degeneration, a leading cause of vision loss, our findings inform organoid design for potential therapeutic applications.

## Introduction

Visual acuity varies dramatically across the animal kingdom (*1*). The high acuity of humans and primates is enabled by the foveola, a specialized region of the central macula (**Fig. 1A-C**). The human retina contains three subtypes of cones: S (expressing short-wavelength, blue opsin), M (expressing medium-wavelength, green opsin), and L (expressing long-wavelength, red opsin) (*2*). The foveola is characterized by a recessed pit containing a high density of M/L cones and the absence of S cones, rods, and vasculature (*3–7*). Whereas the macaque foveola contains a mix of M/L and S cone subtypes, the human foveola is uniquely comprised of exclusively M/L cones (**Fig. 1C**) (*8, 9*). Here, we examined human fetal retinas and retinal organoids to determine mechanisms that generate the human-specific pattern of cone subtypes in the foveola.

**Figure 1.**
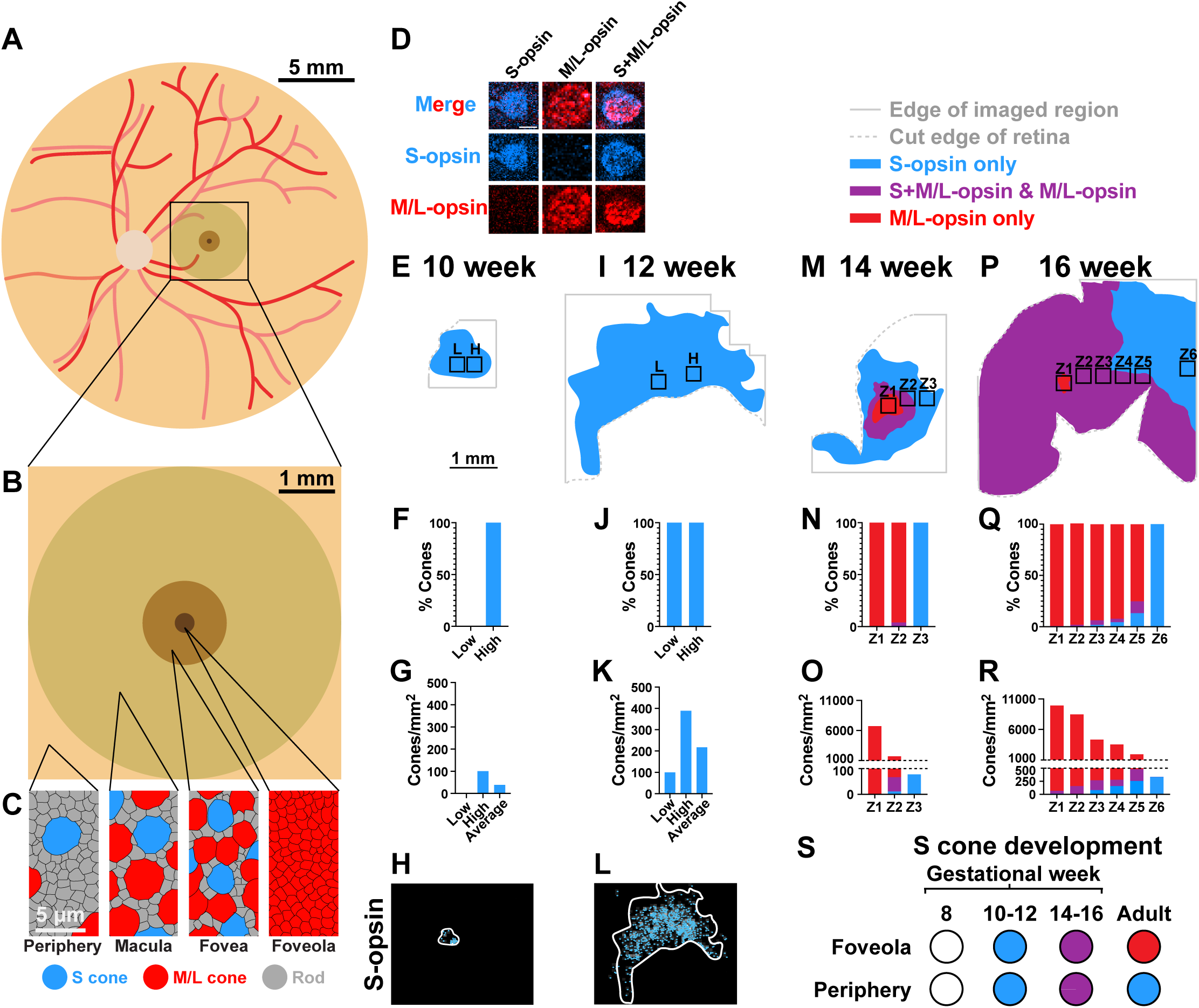
Expression of S-opsin and M/L-opsin during human retinal development. **(A-C)** Adult human retina and photoreceptor pattern schematic. **(A)** Human retina with boxed region representing the macula. Red lines = major arteries, pale lines = major veins, white circle = optic disc. **(B)** Human macula with macular subsections. **(C)** Photoreceptor composition of retinal regions. Grey = rods, blue = S cones, and red = M/L cones. From left to right: the periphery, the macula, the fovea, and the foveola. **(D)** Representative images of S-opsin-expressing, M/L-opsin-expressing, and S+M/L-opsin-expressing cones from a retina at 14 weeks of gestation. Scale bar = 5 μm. **(E, I, M, P)** Representation of cone patterning. Blue region = S-opsin-expressing cones; purple region = M/L-opsin and S+M/L co-expressing cones, and red region = M/L-opsin-expressing cones. Solid lines indicate end of imaged region. Dotted lines indicate cuts in the tissue. Scale bar = 1 mm. **(E)** 10 weeks of gestation, **(I)** 12 weeks of gestation, **(M)** 14 weeks of gestation, **(P)** 16 weeks of gestation. L, H, and Z1-6 indicate quantified zones. **(F, J, N, Q)** Cone subtype composition by % with S-opsin-expressing in blue, S+M/L opsin co-expressing in purple, and M/L-opsin-expressing in red. For the 10- and 12-week retinas, “Low” indicates the region with the lowest S cone density and “High” indicates the region with the highest S cone density. For the 14- and 16-week retinas, Z1-6 indicate quantified zones. **(G, K, O, R)** Cone subtype composition by density with S-opsin-expressing in blue, S+M/L opsin-co-expressing in purple, and M/L-opsin-expressing in red. For the 10- and 12-week retinas, “Low” indicates the region with the lowest S cone density, “High” indicates the region with the highest S cone density, and “Average” indicates the average S cone density across the entire region. For the 14- and 16-week retinas, Z1-6 indicate quantified zones. **(H, L)** Heatmaps overlaid with the locations of each S cone observed at **(H)** 10 weeks of gestation and **(L)** 12 weeks of gestation. **(S)** Summary of observations of S cone development in the presumptive foveola and periphery.

In the current model, cone subtype specification occurs in a two-step process. First, a decision occurs between S and M/L cone fates. If the M/L fate is chosen, a decision is made between M and L fates (*10–13*). As the adult human foveola is composed of both M and L cones and lacks S cones, we focused on the initial decision between S and M/L fates.

In the developing human retina, cone subtype specification initiates in the center and expands to the periphery. S-opsin-expressing cones are first observed in the central retina at fetal week 10. The region containing S-opsin-expressing cones spreads in a wave of differentiation until the entire retina contains S-opsin-expressing cones by fetal week 22 (*14, 15*). M/L-opsin-expressing cones are first observed in the central retina at fetal week 14. M/L cone subtype specification expands to the periphery until the entire retina contains M/L-opsin-expressing cones postnatally (*14*). Though the presumptive foveola contains S-opsin-expressing cones during fetal development (*15*), the adult foveola exclusively contains M/L cones (*4–7*). Based on the observation that the foveola undergoes an increase in cone density in the absence of local cell division or cell death, Diaz-Araya and Provis proposed that M/L cones migrate into the foveola, pushing S cones out (*16*). The absence of an experimentally amenable human model system has limited the ability to directly test this hypothesis and more broadly investigate the mechanisms controlling foveolar cone subtype patterning.

Co-expression of S- and M/L-opsins is observed in the fovea and at the edge of the wave of M/L-opsin expression during fetal eye development (*15, 17*), suggesting potential plasticity of cone subtype fates. Here we test an alternative hypothesis that patterning of the M/L cone-rich foveola is generated by both limited S cone specification and direct conversion of sparse S cones into M/L cones.

While the foveola is unique to primates among mammals, some bird species have analogous high-acuity areas (HAA). Foundational studies in chicken found that CYP26A1, an RA-degrading enzyme, is expressed in the developing HAA. Increasing RA disrupts HAA development (*18*), showing that low levels of RA signaling are critical for the patterning of HAAs. Expression of CYP26A1 in the central retina is conserved in the developing and adult human retina (*18–22*), suggesting a role for RA degradation in cone subtype specification in the human foveola.

We previously used human retinal organoids to show that TH signaling promotes M/L cone specification while repressing S cone specification (*23*). Retinal organoids recapitulate the temporal specification and morphologies of cone subtypes (*23–26*). Our study suggested that dynamic expression of TH-degrading and -activating proteins ensures low TH signaling early to allow specification of S cones and high signaling late to produce M/L cones. Notably, expression of DIO2, an enzyme that converts circulating thyroid hormone (T4) to active thyroid hormone (T3), increases during human fetal retina and organoid development, consistent with both high TH signaling and specification of M/L cones later in development (*23, 27*). Whereas *DIO2* expression decreases in the peripheral retina by adulthood, expression of *DIO2* is maintained at high levels in the central retina (*20, 22*), suggesting that continuous *DIO2* expression and sustained TH signaling promote and maintain the exclusive presence of M/L cones in the foveola.

In this study, we examined cone opsin expression in human fetal retinas and found that, in contrast to the peripheral retina, S-opsin-expressing cones are observed sparsely in the central retina early in development, suggesting active mechanisms that limit their local specification. Later in fetal development, cones in the central retina either express M/L-opsin exclusively or co-express S-opsin and M/L-opsin. By adulthood, this region contains only M/L cones, suggesting that the initial sparse S-opsin-expressing cones may directly convert into M/L cones. Based on the enriched expression of *CYP26A1* and *DIO2* in the foveola, we tested the roles of RA and TH signaling on cone subtype specification in retinal organoids. Our experiments suggest that RA is degraded to promote the specification of M/L cones and limit the generation of S cones in the foveola, and that sustained TH signaling is sufficient to convert S cones to the M/L cone fate, indicating plasticity of cone subtypes even in late developmental stages. Our data suggest a two-step process for cone subtype patterning in the human foveola in which generation of S-cones is initially restricted, followed by a direct conversion of S cones into M/L cones.

## Results

### Cone subtype patterning in the developing foveola

We examined opsin expression in human fetal retinas to describe the developmental dynamics of foveolar cone subtype pattering. Previous studies characterized S-opsin and M/L-opsin expression during development and identified a population of co-expressing cells (*8, 15, 17*). To understand the patterning of cone subtypes, we imaged opsin expression in whole mount fetal retinas. We examined M/L-opsin with an antibody that detects both M- and L-opsin, due to their extremely high protein sequence homology (*2*). At 8 weeks of gestation, we did not observe expression of cone opsins, consistent with previous observations (*15*). At all later timepoints, we observed cells that expressed either S-opsin exclusively, M/L-opsin exclusively, or co-expressed both S-opsin and M/L-opsin (**Fig. 1D**). For each timepoint, we quantified the number of opsin-expressing cells in 300 µm by 300 µm regions, which is approximately the size of the adult S cone-free foveola (*28–30*).

We examined retinas at 10, 12, 14, and 16 weeks of gestation (**Fig. 1E-R**). At gestational week 10, S-opsin-expressing cones were observed in a 990 µm by 450 µm ellipsoid area encompassing the presumptive foveola. To define the range of S-opsin-expressing cone densities, we quantified the regions within this area that contained the most and least cones/mm^2^. The lowest density region contained 0 cones/mm^2^ and the highest density region contained 100 cones/mm^2^ (**Fig. 1E-H, “L” and “H”; Fig. S1A**). The average cone density of the entire region of S-opsin-expressing cones was 38 cones/mm^2^ (**Fig. 1G**). As the overall density of S-opsin-expressing cones was low at this timepoint, we could not determine if the region lacking S-opsin-expressing cones was the presumptive foveola or the result of random variation in the low-density cell pattern.

At gestational week 12, S-opsin-expressing cones covered a region of 1940 μm by 1530 μm, encompassing the presumptive foveola. The lowest density region contained 100 cones/mm^2^ and the highest density region contained 389 cones/mm^2^ (**Fig. 1I-L, “L” and “H”; Fig. S1B**). The average cone density of the entire region of S-opsin-expressing cones was 218 cones/mm^2^ (**Fig. 1K**). The observation of S-opsin-expressing cones throughout all regions suggested that few S-opsin-expressing cones are present in the presumptive foveola at this developmental timepoint.

At gestational week 14, we observed cells that exclusively expressed S-opsin, exclusively expressed M/L-opsin, or co-expressed S-opsin and M/L-opsin. A central region encompassing the presumptive foveola contained cones that expressed M/L-opsin exclusively (**Fig. 1M-O, “Zone 1 (Z1)”; Fig. S1C**). The neighboring region contained a mix of cones that either expressed M/L-opsin exclusively, expressed S-opsin exclusively, or co-expressed S-opsin and M/L-opsin (**Fig. 1M-O, “Z2”; Fig. S1C**). The most peripheral regions only contained cones that exclusively expressed S-opsin (**Fig. 1M-O, “Z3”; Fig. S1C**).

At gestational week 16, we found that the central region encompassing the presumptive foveola mostly contained cones that exclusively expressed M/L-opsin, with some cones that co-expressed S-opsin and M/L-opsin and very few cones that exclusively expressed S-opsin (**Fig. 1P-R, “Z1”; Fig. S1D**). In the four neighboring regions, most cones expressed M/L-opsin exclusively. A proportion of cones expressed S-opsin exclusively or co-expressed S- and M/L-opsin. The proportion of S-opsin-expressing cones increased with the distance from the center (**Fig. 1P-R, “Z2”-“Z5”; Fig. S1D**). A distal region starting at 2.3 mm from the presumptive foveola contained only S-opsin-expressing cones (**Fig. 1P-R, “Z6”; Fig. S1D**). The presence of S-opsin-expressing cones in the central region suggested that small numbers of S-opsin-expressing cones are present in the presumptive foveola, but that most, if not all, S-opsin-expressing cones co-express M/L-opsin during development.

We compared our findings in fetal human retinas to the distribution of cone subtypes in an adult human retina. We quantified cone subtype ratios at various regions beginning near the foveola and moving toward the peripheral retina (**Fig. S1E-H**). Due to the whole mount imaging approach, cone subtype patterning in the foveola was not detectable. At central regions, the retina contained mostly M/L cones (**Fig. S1E, F, I, J**). Towards the periphery, S cones increased in proportion and density (**Fig. S1E, G-J**). Cones that co-expressed S-opsin and M/L-opsin were not observed in the adult retina (**Fig. S1E-J**), suggesting that the co-expression of opsins occurs transiently during fetal retinal development.

Our observations suggest that sparse S cones specified in the foveola initially express S-opsin exclusively, then co-express M/L-opsin, and finally exclusively express M/L-opsin and differentiate as M/L cones (**Fig. 1S**). Outside of the foveola, it appears that S cones initially express S-opsin exclusively, then co-express S-opsin and M/L-opsin, but ultimately revert to exclusively expressing S-opsin (**Fig. 1S**).

Our results are consistent with previous observations of cone subtypes in the central retina during fetal development (*14, 15*). Our experiments validated the presence of S-opsin-expressing cells in the presumptive foveola and assessed the spatial distribution of cones that co-express S-opsin and M/L-opsin. Apoptosis is not readily observed in photoreceptors in the fetal retina, suggesting that the decrease in co-expressing cones is not due to cell death (*17*). Based on these observations, we hypothesized that S cone specification is limited in the foveola and that these sparse S cones are later converted into M/L cones.

### Organoids contain cones that co-express S-opsin and M/L-opsin

We next examined opsin expression in human retinal organoids. To internally control for batch effects and limit variability, organoids were differentiated at the same time for each experiment (**Fig. S2**). Organoids at day 80 contained cones that exclusively expressed S-opsin (**Fig. S3A, S3D-E**). Organoids at day 100 contained cones that exclusively expressed S-opsin or co-expressed S-opsin and M/L-opsin (**Fig. S3D-E**). Organoids at day 110 and day 130 contained cones that exclusively expressed S-opsin, exclusively expressed M/L-opsin, or co-expressed S-opsin and M/L-opsin (**Fig. S3B, S3D-E**). Organoids on days 152, 170, and 190 contained cones that exclusively expressed S-opsin or M/L-opsin (**Fig. S3C-E**). These data suggested two nonexclusive mechanisms of cone subtype development in organoids: (1) S cones express S-opsin exclusively, then co-express S-opsin and M/L-opsin, and finally, revert to expressing S-opsin exclusively, or (2) S cones express S-opsin exclusively, then co-express S-opsin and M/L-opsin, and convert to M/L cones. Both possibilities are consistent with our observations of cone development in human fetal retinas.

### CYP26A1 and low RA promote M/L cone fate and suppress S cone fate

We next sought to identify a mechanism that limits the generation of S-opsin-expressing cones in the foveola. The RA-degrading enzyme CYP26A1 is expressed in the central retina (*18–22*). Low RA signaling is required to pattern the HAA in chicken (*18*). Based on these observations, we hypothesized that CYP26A1-mediated degradation of RA promotes M/L cone fate and suppresses S cone fate.

To investigate the role of CYP26A1 in cone subtype specification, we generated a *CYP26A1* null mutation in human embryonic stem cells (**Fig. S3F-G**). We grew wildtype and *CYP26A1* null mutant retinal organoids in parallel and examined cone subtypes at day 204. Whereas wildtype organoids displayed 98% M/L cones and 2% S cones, *CYP26A1* null mutant organoids displayed 77% M/L cones and 23% S cones (**Fig. 2A-C**), suggesting that CYP26A1 promotes M/L cone and suppresses S cone specification. Wildtype organoids had a higher density of total cones compared to *CYP26A1* null mutants (**Fig. 2D**), suggesting that CYP26A1 likely also promotes cone generation, consistent with the high cone density in the foveola.

**Figure 2.**
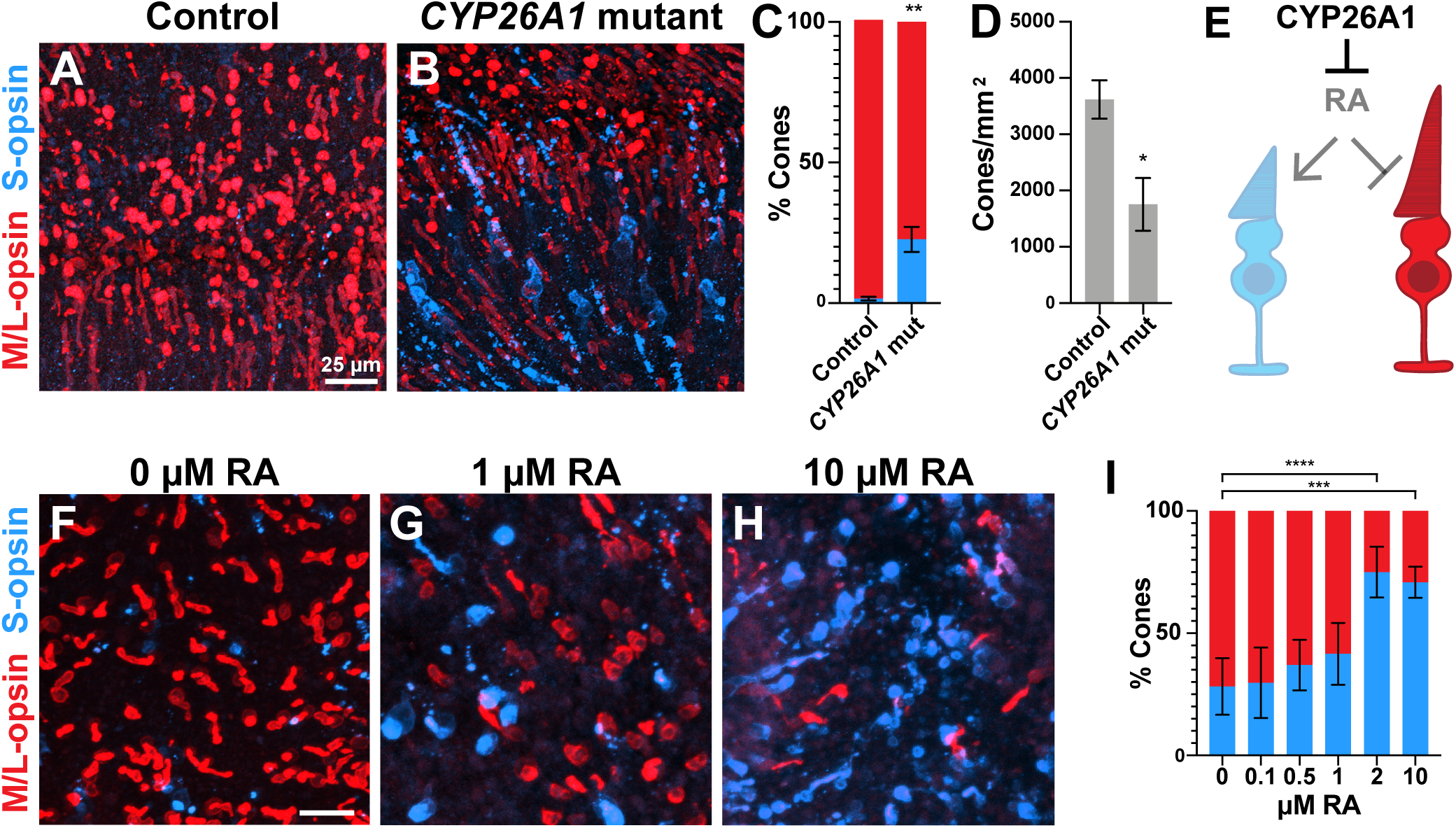
CYP26A1 and low RA limit S and promote M/L cone fate. (**A-B**) Day 204 organoids stained for S-opsin in blue and M/L-opsin in red. Scale bar = 25 μm. **(A)** Control retinal organoid. **(B)** *CYP26A1* mutant retinal organoid. **(C)** Percent cone subtype per genotype (Control, N=3; *CYP26A1* mutant, N=3) with S-opsin in blue and M/L-opsin in red. Unpaired t-test, p=0.0094. **(D)** Cone subtype density per genotype (Control, N=3; *CYP26A1* mutant, N=3). Unpaired t-test, p=0.0327. **(E)** Model of the mechanism through which CYP26A1 promotes M/L and suppresses S cone fate. (**F-H**) Day 213 organoids stained for S-opsin in blue and M/L-opsin in red. Scale bar = 25 μm. **(F)** Organoid treated with 0 μM RA. **(G)** Organoid treated with 1 μM RA. **(H)** Organoid treated with 10 μM RA. **(I)** Percent cone subtype per genotype. S cones in blue and M/L cones in red. One-way ANOVA test with Dunnett correction. N = 5 organoids per condition.

As CYP26A1 degrades RA, we directly tested how levels of RA signaling affect cone subtype specification by growing organoids in different RA concentrations. We found that, as RA concentration increased, the proportion of S cones increased, and the proportion of M/L cones decreased at day 213 (**Fig. 2F-I**). The overall density of all cones was not significantly different across the range of RA concentrations (**Fig. S3H**), suggesting that constant exogenous RA may affect cone development differently than endogenous CYP26A1-mediated regulation. Together, these data suggest that CYP26A1 and low RA suppress S cones and promote M/L cones, consistent with CYP26A1 expression in the central retina and the limited specification of S cones in the foveola (**Fig. 2E**).

### Sustained TH signaling converts S cones into M/L cones

Analyses of fetal retinas suggested that S-opsin-expressing cones in the foveola convert to M/L cone fate (**Fig. 1E-R**) (*15, 17*). We previously showed that TH signaling is necessary and sufficient to promote M/L cone fate and suppress S cones (*23*). *DIO2*, which promotes TH signaling, is expressed at high levels in the central retina through adulthood (*20*). Based on this continuous expression of *DIO2*, we hypothesized that sustained TH signaling converts S cones into M/L cones.

We previously grew human retinal organoids in high active T3 media conditions throughout their development from day 22 to day 200 and observed a high proportion of M/L cones and a low proportion of S cones (*23*). These experiments were conducted in 1 µM RA through day 130 (*23*). For comparison to these observations, we grew organoids similarly in 1 µM RA through day 130 for the next experiments.

To assess how timing and duration of TH signaling influences cone subtype specification, we grew human retinal organoids and varied when we started or ended a pulse of high T3. We defined five key timepoints: before cone generation (day 22), when S-opsin is first detected (day 152), when M/L-opsin is first detected (day 171), after both S and M/L cones are generated (day 190), and when opsin expression is assessed (day 200) (*23*). Organoids for all conditions were grown in parallel from the same differentiation.

We first varied the timing of high T3 pulse termination (“T3 OFF”). Control organoids grown in the absence of supplemental T3 contained a mix of S and M/L with very sparse co-expressing cones (**Fig. 3A, 3E-F**). Organoids grown in high T3 conditions throughout development from days 22-200 were M/L cone-rich (**Fig. 3B, 3E-F**) consistent with our previous observations (*23*). Organoids grown in high T3 conditions beyond S and M/L cone generation from days 22-190 similarly were M/L cone-rich (**Fig. 3E**). Organoids grown in high T3 beyond S cone generation but before M/L cone generation (days 22-171) contained a mix of cones that exclusively expressed M/L-opsin, co-expressed S-opsin and M/L-opsin, or exclusively expressed S-opsin (**Fig. 3E**). Similarly, organoids grown in high T3 until the initiation of S cone generation (days 22-152) contained a mix of cones that exclusively expressed M/L-opsin, co-expressed S-opsin and M/L-opsin, or exclusively expressed S-opsin (**Fig. 3C, 3E**). Adding T3 for extended time windows (day 22-200, 22-190) yielded M/L cone-rich organoids, suggesting that sustained T3 promotes M/L cone and suppresses S cone specification. Adding T3 for shorter periods (day 22-171, day 22-152) generated organoids with cones that co-expressed S-opsin and M/L-opsin, suggesting that a shorter pulse of T3 promotes S cones to take on M/L cone features.

**Figure 3.**
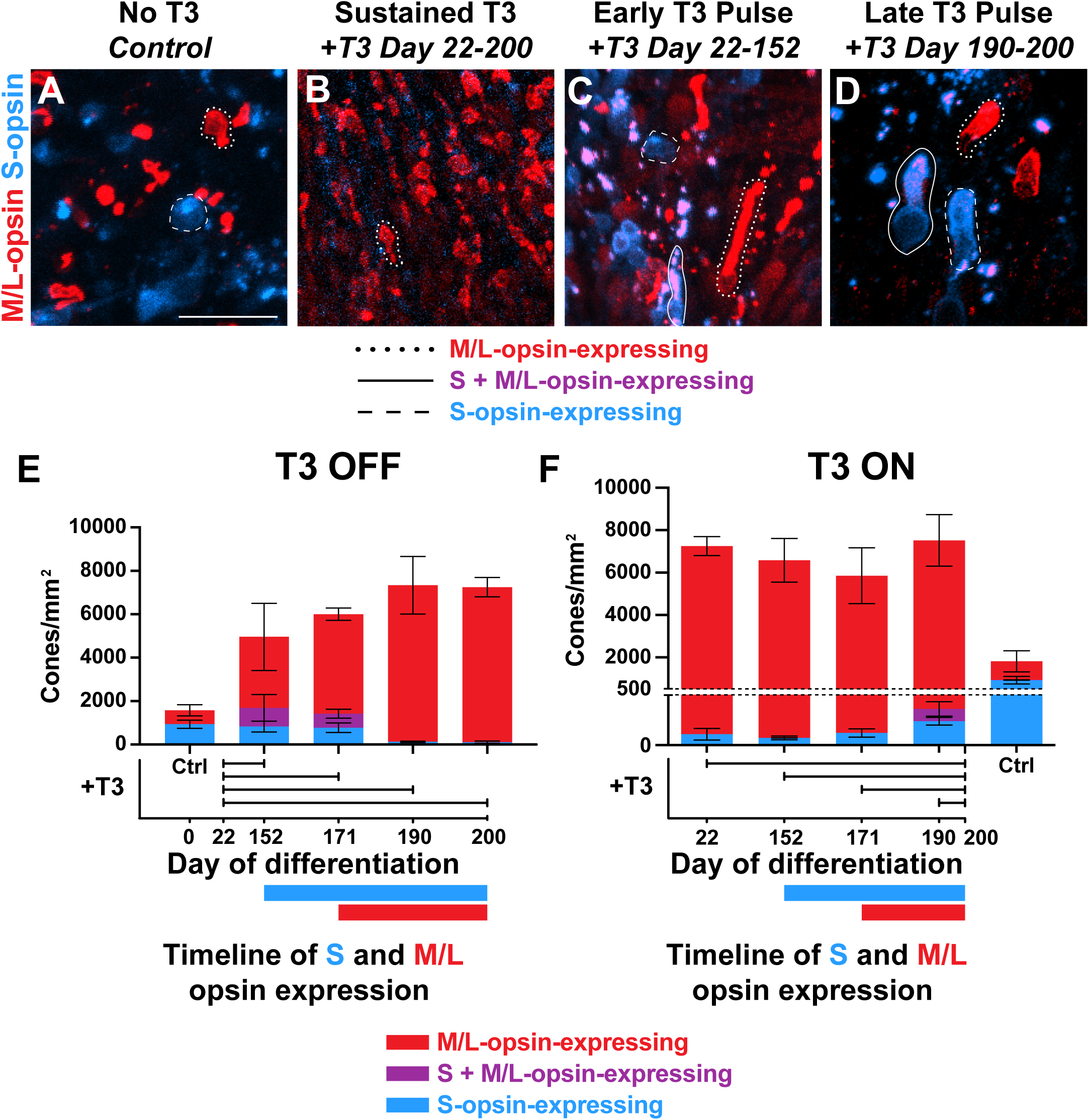
Addition of T3 decreases S-opsin-expressing cells and increases S+M/L-opsin co-expressing and M/L-opsin-expressing cells. **(A-D)** Day 200 organoids. Blue = S-opsin, red = M/L-opsin. Dotted line = M/L-opsin-expressing, solid line = S + M/L-opsin-expressing, dashed line = S-opsin-expressing. Scale bar = 25 μm. **(A)** Control (no addition of T3). **(B)** Addition of 20 nM T3 on days 22-200. **(C)** Addition of 20 nM T3 on days 22-152. **(D)** Addition of 20 nM T3 on days 190-200. **(E-F)** Cone subtype composition by density. Red = M/L-opsin-expressing, purple = S + M/L-opsin-expressing, and blue = S-opsin-expressing. Organoids for these experiments were generated in the same differentiation and grown in parallel. **(E)** T3 OFF. Removal of 20 nM T3 at progressively later timepoints in retinal organoid differentiation. (N=3 for control and *+T3 day 22-200*, N=4 for *+T3 day 152-200*, *+T3 day 171-200*, and *+T3 day 190-200*). **(F)** T3 ON. Addition of 20 nM T3 at progressively later timepoints in retinal organoid differentiation. (N=3 for control, *+T3 day 22-152*, *+T3 day 22-190*, and *+T3 day 22-200*. N=4 for *+T3 day 22-172*).

We next varied the time at which organoids were placed into high T3 conditions (“T3 ON”). Organoids grown in high T3 starting at the initiation of S cone generation (days 152-200) were M/L cone-rich (**Fig. 3F**), suggesting that initiating and sustaining high T3 before S cone generation promotes the specification of M/L cones and suppression of S cones. Organoids grown in high T3 after the onset of S cone generation but before the generation of M/L cones (days 171-200) were M/L cone-rich (**Fig. 3F**), suggesting that T3 is sufficient to promote M/L cones and convert S cones into M/L cones after S cone generation has begun. Organoids grown in high T3 conditions after the generation of S and M/L cones from days 190-200 contained a mix of cones that exclusively expressed M/L-opsin, co-expressed S-opsin and M/L-opsin, or exclusively expressed S-opsin (**Fig. 3D, 3F**), suggesting that this short pulse of T3 induces M/L-opsin expression in S cones, but T3 signaling is not sustained long enough to fully convert all S cones to the M/L cone fate. Together, these data suggest that sustained T3 promotes the generation of M/L cones and the conversion of S cones into M/L cones.

### scRNA-seq shows that T3 regulates cone subtype specification

To extend our analysis of cone subtype changes upon addition of T3, we performed single-cell RNA-sequencing (scRNA-seq) on organoids at day 200 in three conditions: *control* (no T3), *+T3 10-day* (high T3 after the generation of cones; days 190-200), and *+T3 180-day* (high T3 throughout development; days 22-200). These organoids were differentiated in parallel with the organoids analyzed in the previous paragraph and in **Fig 3**. Across all conditions, we recovered transcriptomes of all major retinal cell classes except retinal ganglion cells (**Fig. 4A, S4A-E**), which are lost earlier in organoid development (*31*). To evaluate photoreceptor subtypes, we re-clustered photoreceptor precursors, cones, and rods (**Fig. 4B, S4F-J**).

**Figure 4.**
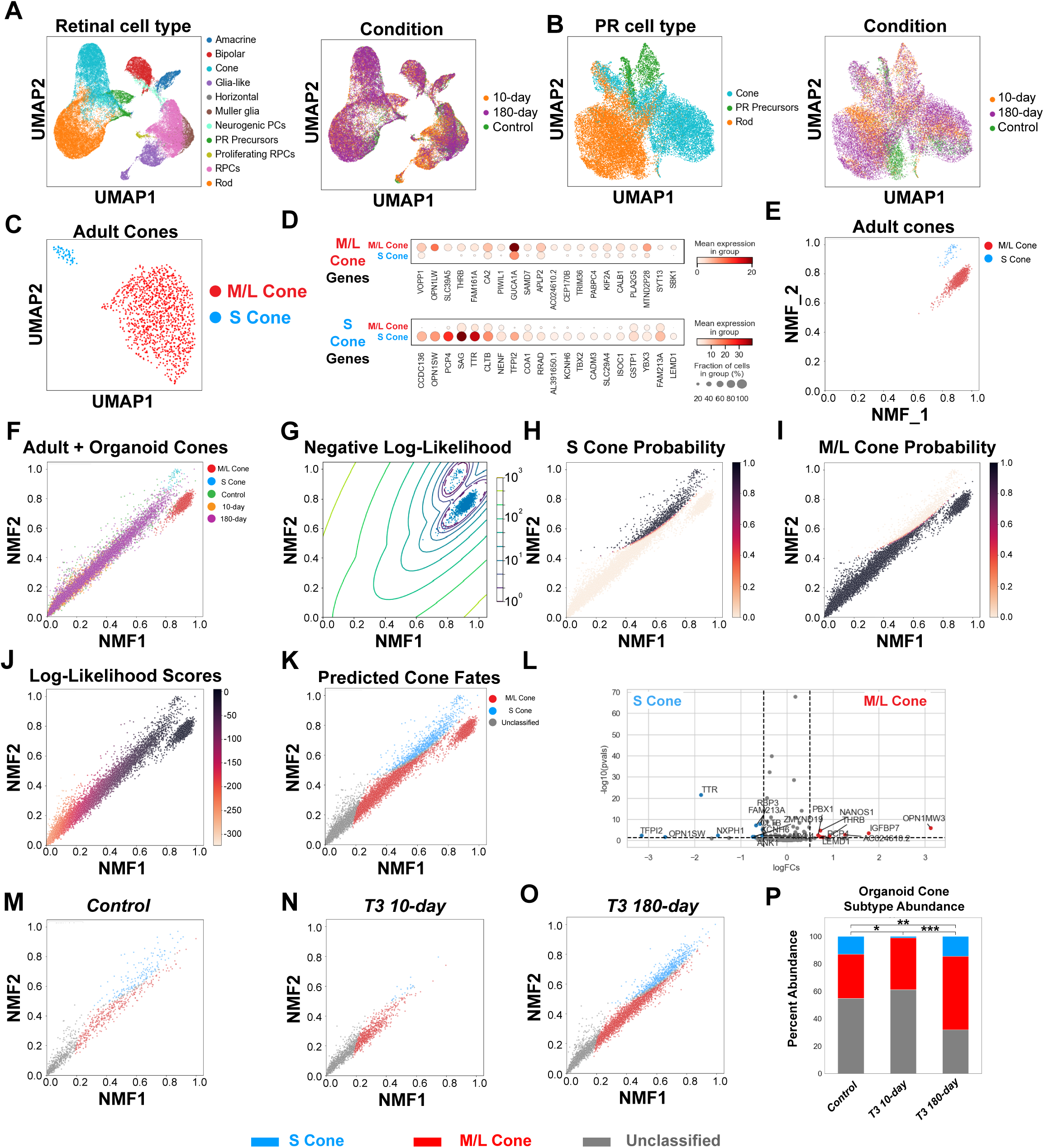
Single-Cell RNA-sequencing analysis of T3-treated organoids. **(A)** Uniform Manifold Approximation and Projection (UMAP) embedding of day 200 organoid cells across three T3 conditions. Left = colored by cell type, right = T3 Condition. **(B)** UMAP embedding of photoreceptor precursors, rods, and cones. Left = colored by cell type, right = T3 Condition. **(C)** UMAP embedding of adult M/L and S cones. Original data from (*19*). **(D)** Top 40 differentially expressed genes between M/L Cones (top) and S cones (bottom). **(E-F)** Embedding of adult M/L and S cones (**E**), and day 200 organoid cones **(F)** onto two NMF components generated from 200 differentially expressed genes. **(G)** Density estimation from adult M/L and S cones in the GMM. **(H-I)** Probabilities of adult and organoid cones assigned as S cones **(H)** or M/L cones **(I)**. **(J)** Log-likelihood scores of adult and organoid cones. **(K)** GMM predicted subtype fates of adult and organoid cones. A cone was assigned a fate if its log-likelihood score exceeded -160 and if its probability for either the S or M/L fate was greater than 0.8. **(L)** Volcano plot of differentially expressed genes between pseudobulked S and M/L cones from organoid cones. LogFC is log2 fold change. (**M-O)** GMM-predicted subtype fates of organoid cones. **(M)** Control. **(N)** *+T3 10-day*. **(O)** *+T3 180-day*. **(P)** Relative abundance of cone subtypes across T3 conditions. Chi square test for proportions. * indicates p<5×10^-3^; ** indicates p< 5×10^-5^; *** indicates p<5×10^-9^.

As rod transcripts are released during cell dissociation and observed in other retinal cell types during scRNA-seq analysis (*20, 32, 33*), we sought to remove cones with a high quantity of rod gene transcripts. We utilized Nonnegative Matrix Factorization (NMF) (*34, 35*), an unsupervised machine learning method, and found gene expression patterns associated with rod genes. We removed cones enriched for these rod gene patterns **(Fig. S5A-C)**.

S and M/L cone subtype clusters were not directly identifiable within the cone cluster (**Fig. 4B**), likely due in part to low opsin mRNA reads. Since opsin mRNA localizes to inner segments (*36*), which break during cell dissociation, this may have contributed to loss of opsin mRNA in these samples. To resolve cone subtypes, we compared cones from organoids to cone subtypes from the adult peripheral retina. Using a published scRNA-seq dataset from the adult peripheral retina (*19*), we generated a UMAP that clustered S and M/L cone subtypes (**Fig. 4C**). From the adult peripheral retina dataset, we identified and ordered the top 200 differentially expressed genes between S and M/L cones (100 S-cone enriched, 100 M/L cone-enriched, **Fig. 4D, S5D-G**).

Using the expression profiles of these 200 differentially expressed genes, we applied NMF to identify two discrete gene expression patterns that describe S and M/L cone subtype fates from the adult cone dataset (NMF1 and NMF2, **Fig. 4E**). We then projected organoid cones onto these latent space representations (**Fig. 4F**). Organoid cones with high projection scores, closer to the adult S and M/L clusters, are likely more mature cones. Organoid cones with low NMF scores are likely more immature cones, S cones converting to the M/L fate, or cones that poorly recapitulate the adult S or M/L fate. As organoid cones are asynchronously developing, we observed a range of cone subtype states (**Fig. 4F**). To quantify S and M/L subtype fates for organoid cones from this projection, we employed a Gaussian Mixture Model (GMM), a method that enables predictions of cluster identities based on training data (*37*). The GMM was trained on the two dimensional latent space representations of S and M/L cones from the adult retina dataset, generating density estimations for each adult S and M/L cone cluster (**Fig. 4G**). Applying these density estimations, organoid cones were classified as specific cone subtypes if their projection into this space exceeded the 80% confidence interval for either the S or M/L cone (**Fig. 4H-I**). Additionally, cones were classified as subtypes if their Log-Likelihood score exceeded -160 (**Fig. 4J**). Based on these criteria, cells were classified as S or M/L cone subtypes (**Fig. 4K**). Cells failing to meet these criteria were deemed as “unclassified” (**Fig. 4K**).

To examine differential gene expression between organoid cones classified as S or M/L subtypes, we performed a pseudobulk analysis. Notably, in aggregate, S-opsin expression was enriched in cones classified as S subtype (**Fig. 4L**). *M-opsin* expression (*OPN1MW* and *OPN1MW3*) was enriched in cones classified as M/L subtype (**Fig. 4L**). Detection of *M-opsin*, but not *L-opsin*, is consistent with our previous observations that addition of RA in organoids promotes M and suppresses L cone fate (*38*).

Of the top 20 differentially expressed genes between organoid S and M/L cones, 17 were enriched in the same subtype in adult cones. For example, TTR was enriched in both organoid and adult S cones and IGFBP7 was enriched in both organoid and adult M/L cones. Three genes were differentially expressed in the opposite subtype, including MYL4, PCP4, and LEMD1 (**Fig. 4L**).

We next compared populations of cone subtypes from *control*, *+T3 10-day*, and *+T3 180-day* organoids. S, M/L, and unclassified cone subtypes were identified in all three conditions (**Fig. 4M-P**). Compared to *control* organoids, *+T3 10-day* organoids displayed an increase in the proportion of unclassified cones and a decrease in S cones (**Fig. 4P**), consistent with S cones converting into M/L fate (**Fig. 3D, 3F**). *+T3 180-day* organoids displayed an increase in the proportion of M/L cones (**Fig. 4P**), recapitulating our observations based on opsin protein imaging (**Fig. 3B, 3F**).

Together, our imaging (**Fig. 3**) and scRNA-seq (**Fig. 4**) analysis suggest that S-opsin-expressing cells convert to M/L cone fate in high T3 conditions.

## Discussion

We investigated the patterning of cone subtypes in the human foveola. The foveola initially contains sparse S-opsin-expressing cones. During fetal development, the foveola contains cones that express M/L-opsin only or co-express S-opsin and M/L-opsin. By adulthood, all cones in the foveola exclusively express M/L-opsin (*4–7*). Whereas wild type organoids and low RA conditions resulted in increased M/L cones, *CYP26A1* mutants and high RA conditions yielded organoids with increased S cones. While pulses of T3 yielded organoids with cones that co-expressed S- and M/L-opsin and an increase in unclassified cones, sustained high levels of T3 generated organoids enriched in M/L cones. Together, these data suggest that CYP26A1 degrades RA to limit S cone generation (**Fig. 5A**), and sustained T3 signaling converts sparse S cones into M/L cones (**Fig. 5B**), to generate the M/L cone-rich pattern in the foveola (**Fig. 5C**).

**Figure 5.**
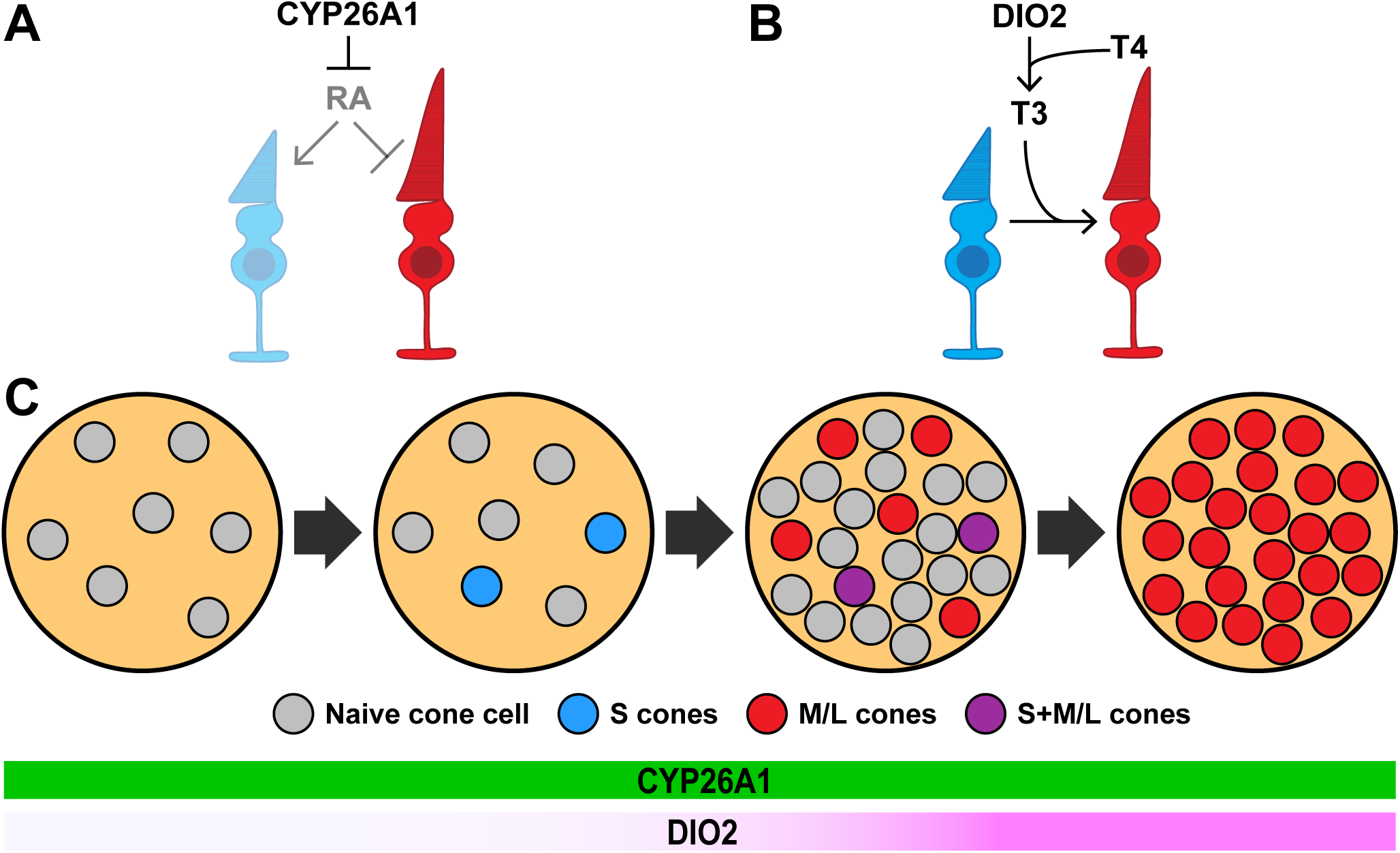
Model for foveolar cone patterning. **(A)** Model: CYP26A1 represses RA to promote M/L cone fate. **(B)** Model: Prolonged DIO2 converts inactive T4 into active T3, converting S cones to M/L cone fate. **(C)** Model of the temporal mechanism of foveolar cone subtype specification in humans.

Our findings for the roles of RA and TH in human cone subtype specification and conversion are consistent with observations from other species. In mice, expression of *CYP26A1* is restricted to a stripe in the central retina by the Vax2 transcription factor. In *Vax2* knockouts, the expression domain of CYP26A1 is expanded, as is the region containing M cones into the predominantly S-cone-rich dorsal retina (*39*), consistent with the role of CYP26A1 in promoting M/L and limiting S cone fate in human retinal organoids. Opsin switching has been observed in several species. In salmonids, TH controls switching of opsin expression from UV opsin to blue opsin during smoltification (*40, 41*).

In flies, ecdysone hormone mediates a blue to green switch in opsin expression in photoreceptors of the Bolwig’s organ during the transition from larval to adult development (*42*). Rat and gerbil retinas also undergo periods of opsin co-expression at the onset of M cone specification, which resolves as the animals mature, though the mechanisms are not understood (*43*). These observations are consistent with our finding that sustained TH signaling converts S cones to M/L cones in humans.

Our studies show that organoids can be used to model and study human-specific regional patterning of the retina. Our studies have important implications for developing therapies for retinal degenerative diseases, including macular degeneration. While current therapies slow disease progression (*44–48*), there is no cure. Transplantation approaches could be improved using retinal organoids with specific ratios of photoreceptors that match the transplanted region. Our work identified genetic and pharmacological manipulations that pattern cone subtypes in organoids similar to the high-acuity foveola. Since the foveola is highly susceptible to impairment in diseases including macular degeneration, our findings can help guide organoid protocols for therapeutic applications.

## Supplemental Materials

## Materials and Methods

### Cell line maintenance

H7 ESC (WA07, WiCell) cells were used for differentiation. Stem cells were maintained in mTeSR™1 (85857, StemCell Technologies) on 1% (v/v) Matrigel-GFR™ (354230, BD Biosciences) coated dishes and grown at 37°C in a HERAcell 150i or 160i 10% CO_2_ and 5% O_2_ incubator (Thermo Fisher Scientific). Cells were passaged every 4-5 days according to confluence as in (*1*). Cells were passaged with Accutase (SCR005, Sigma) for 7-12 minutes and dissociated to single cells. Cells in Accutase were added 1:2 to mTeSR™1 with 5 µM Blebbistatin (Bleb, B0560, Sigma), and pelleted at 150 x g for 5 minutes. Cells were resuspended in mTeSR™1 with 5 µM Blebbistatin, density was quantified via hemocytometer, and cells were plated at 5,000-15,000 cells per well in a prepared 6-well Matrigel coated plate. Cells were fed with mTeSR™1 48 hours after passaging and every subsequent 24 hours until passaged again. To minimize cell stress, no antibiotics were used. Cell lines were tested monthly for mycoplasma using MycoAlert (LT07, Lonza).

#### Cell culture media

##### Stem cell media

mTeSR™1 (85857, StemCell Technologies)

##### E6 supplement

970 µg/mL Insulin (11376497001, Roche), 535 µg/mL holotransferrin (T0665, Sigma), 3.20 mg/mL L-ascorbic acid (A8960, Sigma), 0.7 µg/mL sodium selenite (S5261, Sigma).

##### BE6.2 media

2.5% E6 supplement (above), 2% B27 supplement (50X) minus Vitamin A (12587010, Gibco), 1% Glutamax (35050061, Gibco), 1% NEAA (11140050, Gibco), 1 mM Sodium Pyruvate (11360070, Gibco), and 0.87 mg/mL NaCl in DMEM (11885084, Gibco).

##### LTR (Long-term retina) media

25% F12 (11765062, Gibco), 2% B27 supplement (50X) (17504044, Gibco), 10% heat-inactivated FBS (16140071, Gibco), 1mM Sodium Pyruvate (11360070, Gibco), 1% NEAA (11140050, Gibco), 1% Glutamax (35050061, Gibco), and 1 mM taurine (T-8691, Sigma) in DMEM (11885084, Gibco).

##### Retinoic acid treatment

For organoids, stocks of 10.4 mM retinoic acid (ATRA; R2625, Sigma) were made in DMSO. Retinoic acid was then dissolved in LTR to reach desired working concentration.

##### Thyroid hormone treatment

For organoids, 20 nM T3 (T6397, Sigma) in LTR as used previously in mouse retinal explant culture (*2*) and human retinal organoids (*3*).

#### Organoid differentiation

Organoids were differentiated from H7 WA07 ESCs as described in (*1, 3*) (**Fig. S2**). Briefly, pluripotent stem cells were well-maintained, and only cultures with minimal-to-no spontaneous differentiation were used for aggregation. To aggregate, cells were passaged in Accutase at 37°C for 13 minutes to ensure complete single-cell dissociation. Cells were seeded in 50 uL of mTeSR™1 at 3,000 cells/well into 96-well ultra-low adhesion round-bottom Lipidure coated plates (51011610, NOF). Cells were placed in hypoxic conditions (10% CO2 and 5% O2) for 24 hours to enhance survival. Cells naturally aggregated by gravity over 24 hours.

On day 1, cells were moved to normoxic conditions (5% CO2). On days 1-3, 50 uLs of BE6.2 media containing 3 μM Wnt inhibitor (IWR1e: 681669, EMD Millipore) and 1% (v/v) Matrigel were added to each well. On days 4-9, 100 uL of media were removed from each well, and 100 uL of media were replenished. On days 4-5, BE6.2 media containing 3 μM Wnt inhibitor and 1% Matrigel was added. On days 6-7, BE6.2 media containing 1% Matrigel was added. On days 8-9, BE6.2 media containing 1% Matrigel and 100 nM Smoothened agonist (SAG; 566660, EMD Millipore) was added.

On day 10, aggregates were transferred to 15 mL tubes, rinsed 2-3X in DMEM (11885084, Gibco), and resuspended in BE6.2 with 100 nM SAG in untreated 10 cm polystyrene petri dishes. From this point on, media was changed every other day. Aggregates were monitored daily and manually separated if stuck together or to the bottom of the plate.

On days 13-16, LTR media with 100 nM SAG was utilized during media changes. Between days 11 and 16, retinal vesicles were optionally manually dissected if needed using sharpened tungsten needles. After dissection, cells were transferred into 15 mL tubes and washed 2X with 5 mLs of DMEM.

On days 16-20, cells were maintained in LTR and washed 2X with 5 mLs of DMEM, before being transferred to new plates to wash off dead cells. To increase survival and differentiation, 1 uM all-trans retinoic acid (ATRA; R2625; Sigma) was added to LTR medium from days 20-43 in all conditions. After day 43, retinoic acid was used at varying timings and concentrations depending on experimental conditions. 10 μM Gamma-secretase inhibitor (DAPT; 565770, EMD Millipore) was added to LTR from days 28-42. Organoids were grown at low density (10-20 per 10 cm dish, 2-3 per well in 6 well plate) to reduce aggregation. Organoids were regularly culled if failing to differentiate properly.

#### CRISPR mutations

##### Cloning gRNA plasmids

Plasmids for gRNA transfection were generated using pSpCas9(BB)-P2A-Puro plasmid modified from the pX459_V2.0 plasmid (62988, Addgene) by replacing T2A with a P2A sequence. gRNAs were cloned into the vector following the Zhang lab protocol (*4, 5*).

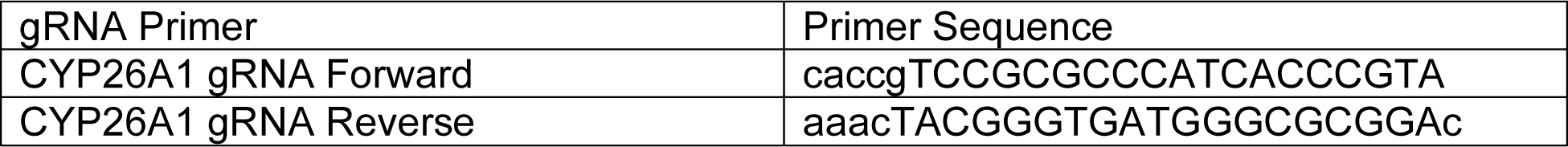

##### Transfection and mutation identification

H7 stem cells were passaged in Accutase at 37°C for 13 minutes to ensure single-cell dissociation. Cells were seeded at 50,000 cells/well in a Matrigel-coated 24-well plate for 24 hours in mTeSR™1 with 5 µM Blebbistatin at hypoxic conditions. After 24 hours, media was removed and fresh mTeSR™1 was added. Cells were transfected with 2.5 uL Lipofectamine Stem Reagent (STEM00015, ThermoFisher Scientific), 200 ng gRNA plasmid PX459v2 containing the gRNA and Cas9-p2a-puromycin-resistance genes in 50 uL of Opti-MEM (31985062, Gibco). Cells were incubated for 24 hours, then media was removed and fresh mTeSR™1 was added. After 24 hours, media was removed and fresh mTeSR™1 along with 0.3-1 ug of puromycin (P8833, Sigma-Aldrich) was added to generate a kill curve. After 24 hours, media was removed and cells were washed 1X with mTeSR™1, then fresh mTeSR™1 was added to the wells. After several days, cells surviving at a puromycin concentration where all control cells had died were passaged at single cell density into a new, Matrigel-coated 6-well plate. Individual colonies were then isolated and mutations were confirmed by PCR sequencing. A gene diagram of the deletion is displayed in **Fig. S3F-G**.

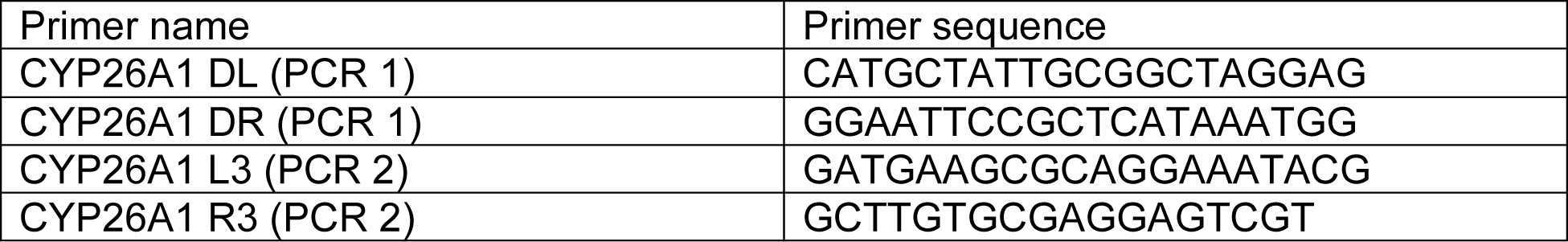

#### Immunohistochemistry

##### Retinal organoids

Retinal organoids were fixed in fresh 4% formaldehyde and 5% sucrose in PBS for 50 minutes. Tissue was rinsed 3X in 5% sucrose in PBS, then incubated at 4°C in 6.75% sucrose in PBS for 30 min, 12.5% sucrose in PBS for 30 min, and 25% sucrose for 2 hours-overnight. Organoids were incubated for 2 hours-overnight at 4°C in blocking solution (0.2-0.3% Triton X100, 2-4% donkey serum in PBS). Organoids were incubated with primary antibodies in blocking solution for 16-36 hours at 4°C. Organoids were washed 3X for 15 minutes in PBS, and then incubated with secondary antibodies in blocking solution overnight at 4°C. Organoids were washed 3X for 10 minutes in PBS. Organoids were optionally incubated in 300 nM DAPI in blocking solution for 10 min and washed 3X for 15 min in PBS. At the end of staining, organoids were mounted for imaging in slow fade (S36940, Thermo Fisher Scientific).

##### Retinas

Human retinas were obtained from the National Disease Research Interchange (NDRI), the University of Maryland Brain and Tissue Bank, and the laboratory of Tom Reh. Human retinal tissue was fixed in 10% formalin within 12 hours post-mortem and stored at 4°C until dissection. Retinas were dissected and whole-mounted, then rinsed 3X in PBS for 20 min, and blocked for 48 hours at 4°C in 0.3% Triton X-100 and 4% donkey serum. Retinas were stained with the same protocol as detailed above for organoids.

#### Antibodies

Primary antibodies were used at the following dilutions: goat anti-SWopsin (1:200 for organoids, 1:500 for human retinas) (sc-14363 Santa Cruz Biotechnology), chick anti-SWopsin (1:200 for organoids, 1:500 for human retinas) (gift from the laboratory of Jeremy Nathans) rabbit anti-LW/MW-opsins (1:200 for organoids, 1:500 for human retinas) (AB5405 Millipore), and mouse anti-Rhodopsin (1:500) (GTX23267 GeneTex). All secondary antibodies were Alexa Fluor-conjugated (1:400) and made in donkey (Molecular Probes).

#### Microscopy and image processing

Fluorescent images were acquired with a Zeiss LSM800 or LSM980 laser scanning confocal microscope. Confocal microscopy was performed with similar settings for laser power, photomultiplier gain and offset, and pinhole diameter. Maximum intensity projections of z-stacks (5–150 optical sections, 1.00 μm step size) were rendered to display all cones captured in a single organoid.

#### Measurements and Quantification

Measurements of cone density were done using ImageJ software. Quantifications and statistics (except for RNA-seq data) were done in GraphPad Prism, with a significance cutoff of 0.01. Statistical tests are listed in figure legends. All error bars represent the SEM. Organoids with < 150 total cones were deemed unhealthy and removed from our analysis.

#### Single-cell RNA sequencing

Retinal organoids were dissociated using Papain Dissociation System (LK003150, Worthington). Briefly, individual organoids were dissociated in a papain + DNase solution for 60 minutes at 37°C with gentle mixing every 5 minutes. The cell suspension mixture was then gently triturated 30 times to break up any remaining tissue chunks before being placed on an ovomucoid discontinuous density gradient to remove cell debris. The dissociated cells were then resuspended with ice-cold PBS containing 0.04% bovine serum albumin (BSA) and 0.5 U/μl RNase inhibitor and were filtered through a 40-μm filter. Cell counts and viability was assessed by Trypan blue staining before loading the cells on a 10x Chromium controller for scRNA sequencing.

#### Single-cell RNA sequencing preprocessing and analysis

Sequencing reads were demultiplexed and aligned to the GRCh38 human reference genome using the Cell Ranger v3.1 *mkfastq* and *count* pipeline (10× Genomics, Pleasanton, CA) with default parameters for scRNA-seq. The resulting cell-by-gene count matrix served as input for downstream analysis.

The count matrix was processed with the Seurat 4.0 R package (*6*). Cells were filtered to exclude those with less than 25% of counts mapped to mitochondrial genes, more than 1000, and fewer than 20000 unique molecular identifiers (UMIs). Doublets were identified and removed using the scDblFinder R package (*7*). Subsequently, cells from individual datasets were merged, normalized and scaled using Seurat’s *SCTransform* function, with the percentage of mitochondrial counts and ribosomal counts used as variables for the vars.to.regress option. Principal components were calculated based on the top 2000 variable features, and Uniform Manifold Approximation and Projection (UMAP) dimension reduction was performed on the top 14 principal components. Clusters were computed using Seurat’s *FindNieghbors* and *FindClusters* functions at a resolution of 1.4. Cell types were identified using a list of known marker genes previously documented (*8–10*). All scripts used for preprocessing can be found at github: https://github.com/csanti88/hussey_t3treatedorganoids_scRNA_2022.

To examine cone subtypes, we used NMF (*sklearn*, n_components = 20, max_iter = 30,000) on the organoid cones. We found an NMF pattern (NMF_8) heavily loaded with rod associated genes (NRL, RHO). We removed cells with high NMF_8 values to remove falsely annotated rods **(Fig. S5A-C)**.

Adult cone subtypes were subset from (*11*). NMF was performed on the top 2000 highly variable genes selected by Pearson residuals, and UMAPs were generated using *scanpy*’s UMAP function with default parameters (**Fig. 4B, S4F-J**) (*11, 12*). We identified differentially expressed (DE) genes between adult S and M/L cones via scanpy’s rank_genes_group tool (n_genes = 100, method = ‘wilcoxon’) (**Fig. 4D, S5E**). To determine the number of genes that explain the differences between adult S and M/L cones, we examined the variance explained by the first principal component of this dataset for subsets containing variable numbers of DE genes (**Fig. S5D**). We reasoned that the top 200 DE genes (100 S cone enriched, 100 M/L cone enriched) were sufficient to explain the differences between adult S and M/L cones while also filtering out noise (**Fig. S5D**). NMF (n_components = 2) were used on the top 200 DE genes. Organoid cones were projected onto these NMF patterns using scProject (**Fig. 4F**). A GMM (sklearn, n_comps = 2) was trained on adult S and M/L cones, and the identities of organoid cones were then classified (**Fig. 4G-K**). Cones with a log-likelihood exceeding - 160 and an 80% confidence interval for the S or M/L fate were annotated a subtype fate, otherwise they were given an ‘unclassified’ cone fate (**Fig. 4H-K**). To ensure the GMM annotations were biologically relevant, we pseudo-bulked organoid S and M/L cones and performed differential gene expression analysis using *decoupler* and *pydeseq2* (**Fig. 4L**). All scripts used for subsetting cones and subsequent analyses can be found on github: https://github.com/BGuy2/HusseyFoveaPatterning.

---

**Supplemental Figure 1.**
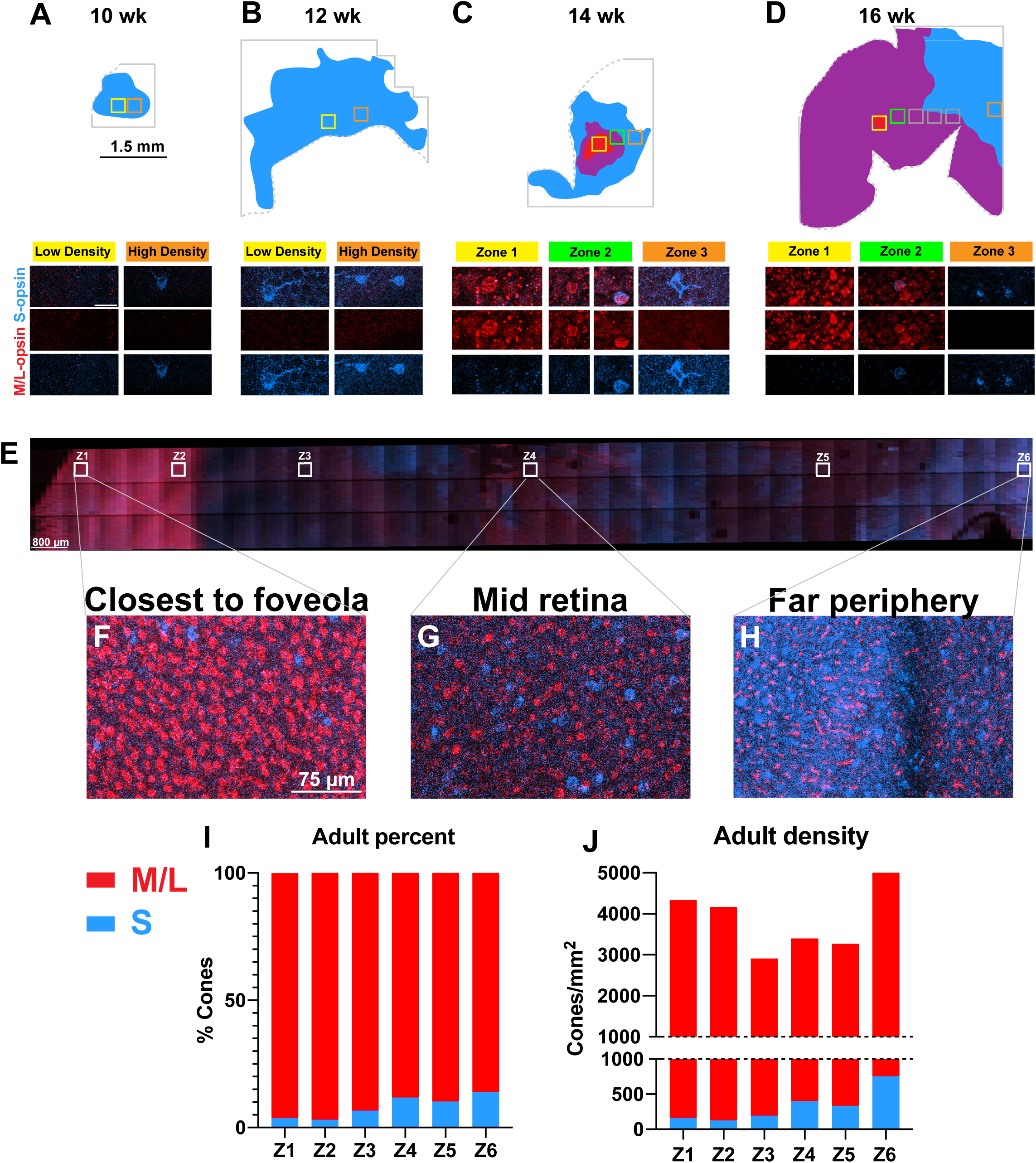
Distribution of cones during development. (**A-D**) Top, representation of cone patterning. Blue region = S-opsin-expressing cones; purple region = M/L-opsin and S+M/L opsin co-expressing cones, and red region = M/L-opsin-expressing cones. Solid lines indicate end of imaged region. Dotted lines indicate cuts in the tissue. Scale bar = 1.5 mm. Solid lines indicate end of imaged region. Dotted lines indicate cuts in the tissue. Bottom, representative images of photoreceptors. **(E)** Adult human retina from near foveola to far periphery. Scale bar = 800 μm. Boxes indicate representative zones quantified in **(I)** and **(J)**. **(F-H)** Representative images of adult human retinal regions **(F)** near foveola, **(G)** in the mid-retina, **(H)** in the far periphery. Scale bar = 75 μm. **(I)** Percent cone subtypes in an adult human retina. Red = M/L-opsin-expressing, blue = S-opsin-expressing. **(J)** Density of cone subtypes in an adult human retina. Red = M/L-opsin-expressing, blue = S-opsin-expressing.

**Supplemental Figure 2.**
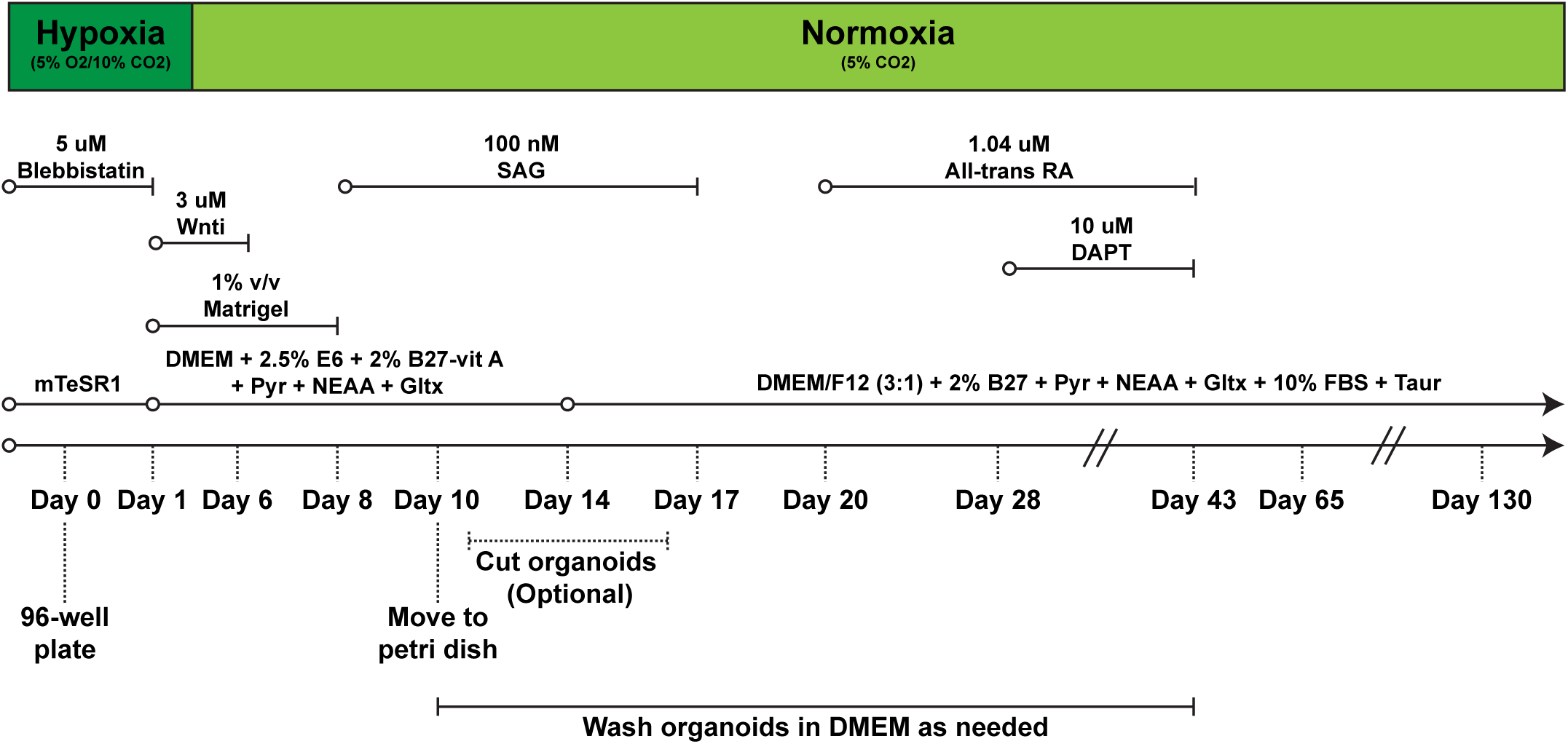
Retinal organoid differentiation protocol. Standard conditions for human retinal organoid differentiations as described in the materials and methods. Method adapted from (*23, 25*).

**Supplemental Figure 3.**
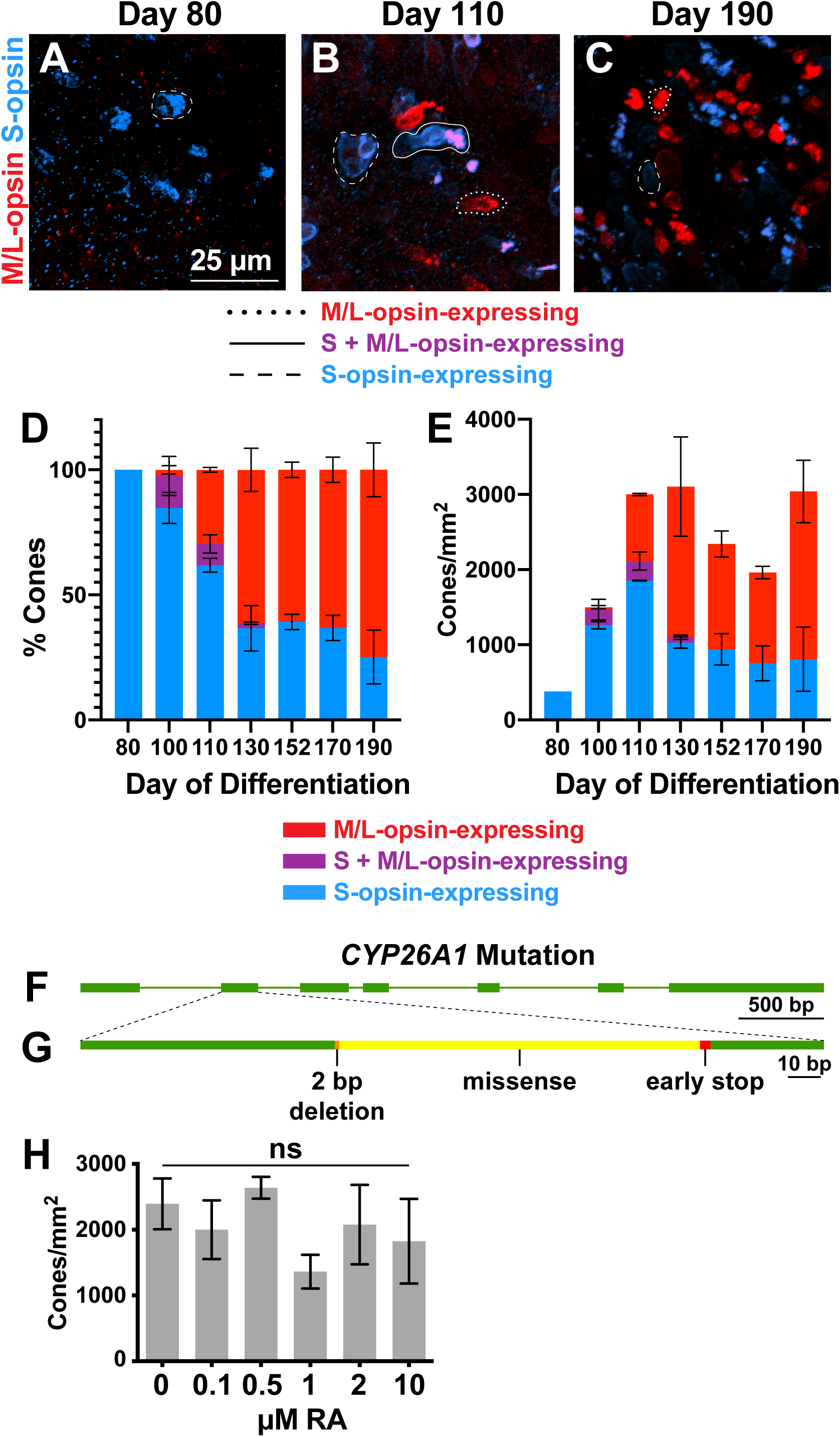
Co-expression of S- and M/L-opsin during organoid development; CYP26A1 mutation; RA and cone densities. **(A-E)** Time course of human retinal organoid cone subtype specification. (N=3 for all timepoints). Scale bar = 25 μm. **(A)** Retinal organoid on day 80. **(B)** Retinal organoid on day 110. **(C)** Retinal organoid on day 190. **(D)** Percent cone subtypes during retinal organoid development. **(E)** Density of cone subtypes during retinal organoid development. **(F-G)** Schematic of the CRISPR mutation generated in the CYP26A1 gene **(F)** Schematic depicting the *CYP26A1* gene. Green squares = exons, Green lines = introns. **(G)** Schematic depicting the nature of the mutation within exon 2. Green = unchanged, orange = deleted bases, yellow = a region of out of frame sequence, and red = early stop codon. (**H**) Total cone density does not change significantly in different RA conditions. Related to **Fig. 2I**.

**Supplemental Figure 4.**
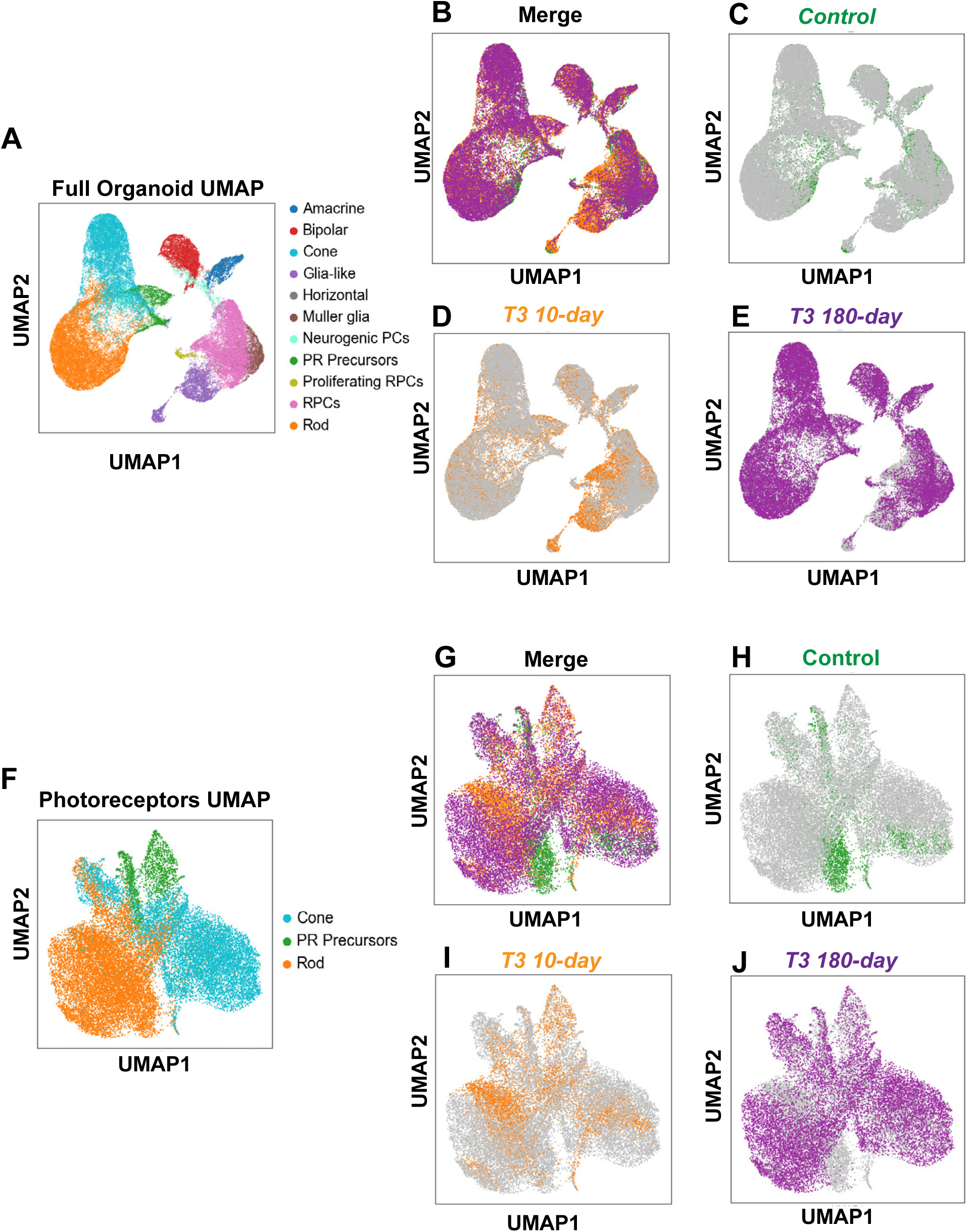
UMAPs of day 200 retinal organoids. **(A)** UMAP embedding of day 200 retinal organoids, colored by cell type. **(B)** UMAP embedding as in **A**, colored by condition. **(C-E)** UMAP embeddings as in **A.** **(C)** *Control*. **(D)** *+T3 10-day*. **(E)** *+T3 180-day*. **(F)** UMAP embedding of photoreceptor cell types from day 200 retinal organoids, colored by cell type. **(G)** UMAP embedding as in **F**, colored by condition. **(H-J)** UMAP embeddings as in **F**. **(H)** *Control*. **(I)** *+T3 10-day*. **(J)** *+T3 180-day*.

**Supplemental Figure 5.**
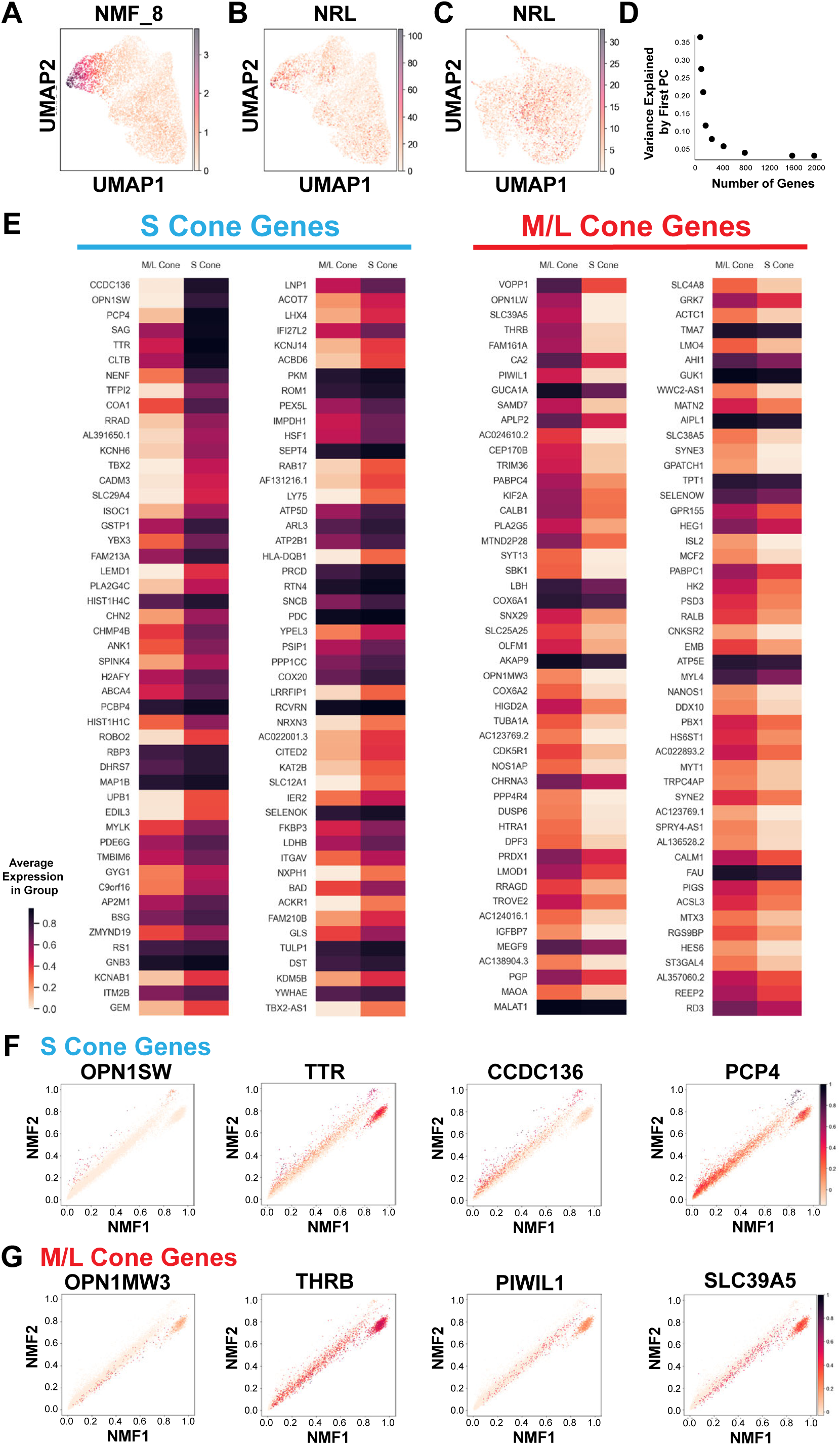
Cone cell filtering and subtype-enriched gene selection for analysis. **(A)** UMAP embedding of day 200 organoid cones, colored by NMF8 score. NMF8 was weighted highly with rod-associated genes. **(B)** UMAP embedding of day 200 organoid cones, colored by NRL (rod-specific transcription factor gene) expression in read counts. **(C)** UMAP embedding of day 200 organoid cones, filtered by NMF8 score to remove rod contaminants, colored as in **(B)**. **(D)** Variance explained by the first principal component across varying numbers of differentially expressed genes in adult S and M/L cones. **(E)** Top 100 differentially expressed genes in adult S (left) and M/L (right) cones. RNA expression shown as a fraction of the gene’s maximum read count from adult cones subtypes. **(F-G)** RNA expression shown as a fraction of the gene’s maximum read count in organoid and adult cones. **(F)** Expression patterns of S-cone associated genes in organoid cones. From left to right: OPN1SW, TTR, CCDC136, and PCP4. **(G)** Expression patterns of M/L-cone associated genes in organoid cones. From left to right: OPN1MW3, THRB, PIWIL1, SLC39A5.

